## Supplemental Figures for "Nicotine enhances the stemness and tumorigenicity in intestinal stem cells via Hippo-YAP/TAZ and Notch signal pathway"

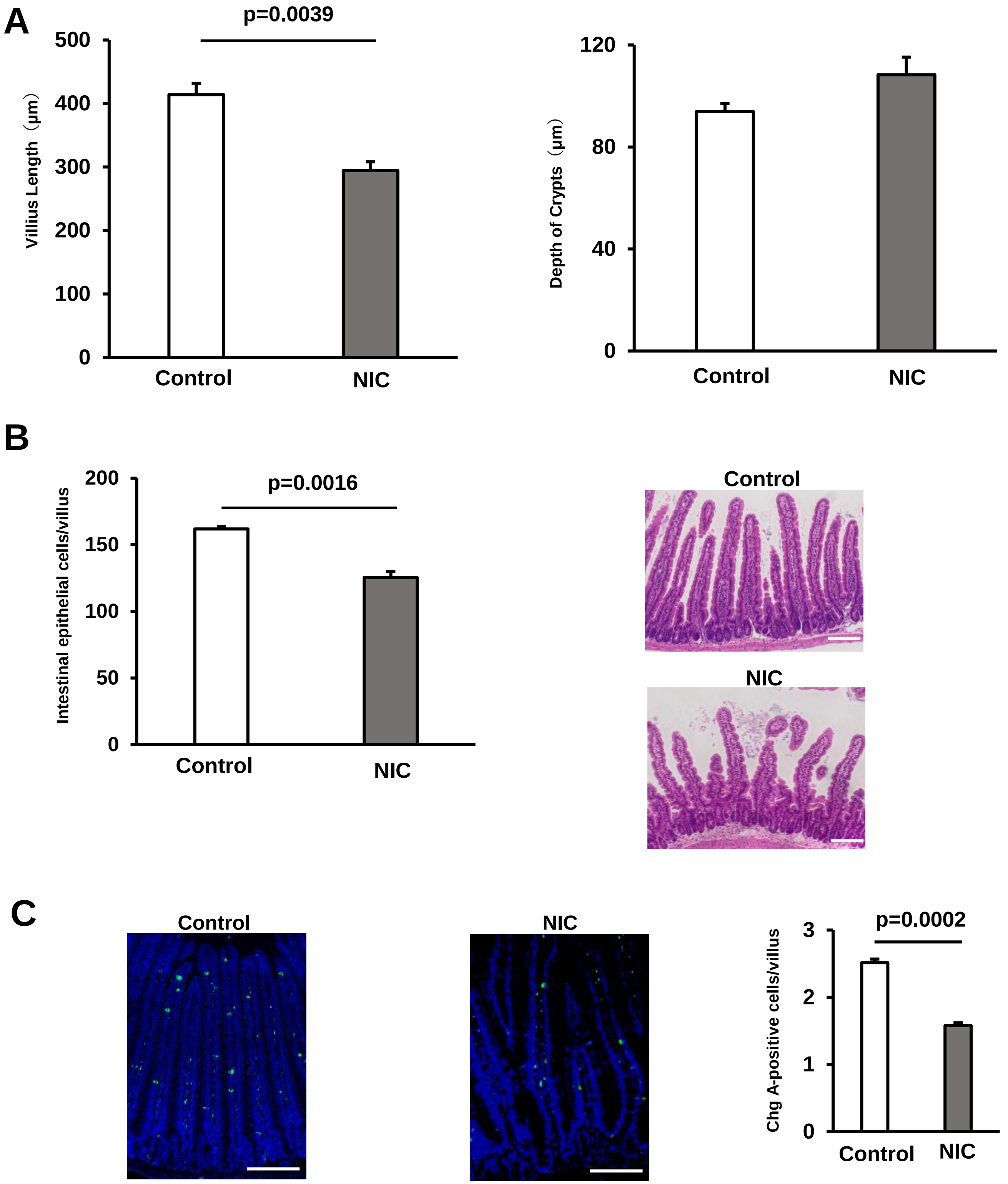


### Figure1 Supplemental

A

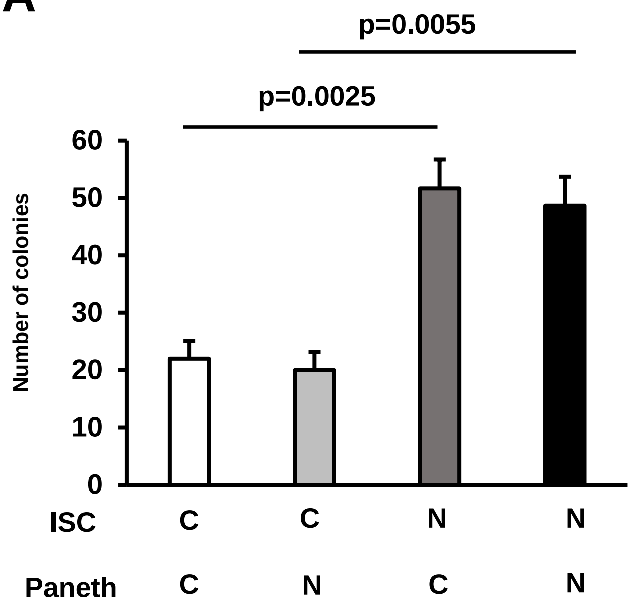


### Figure2 Supplemental

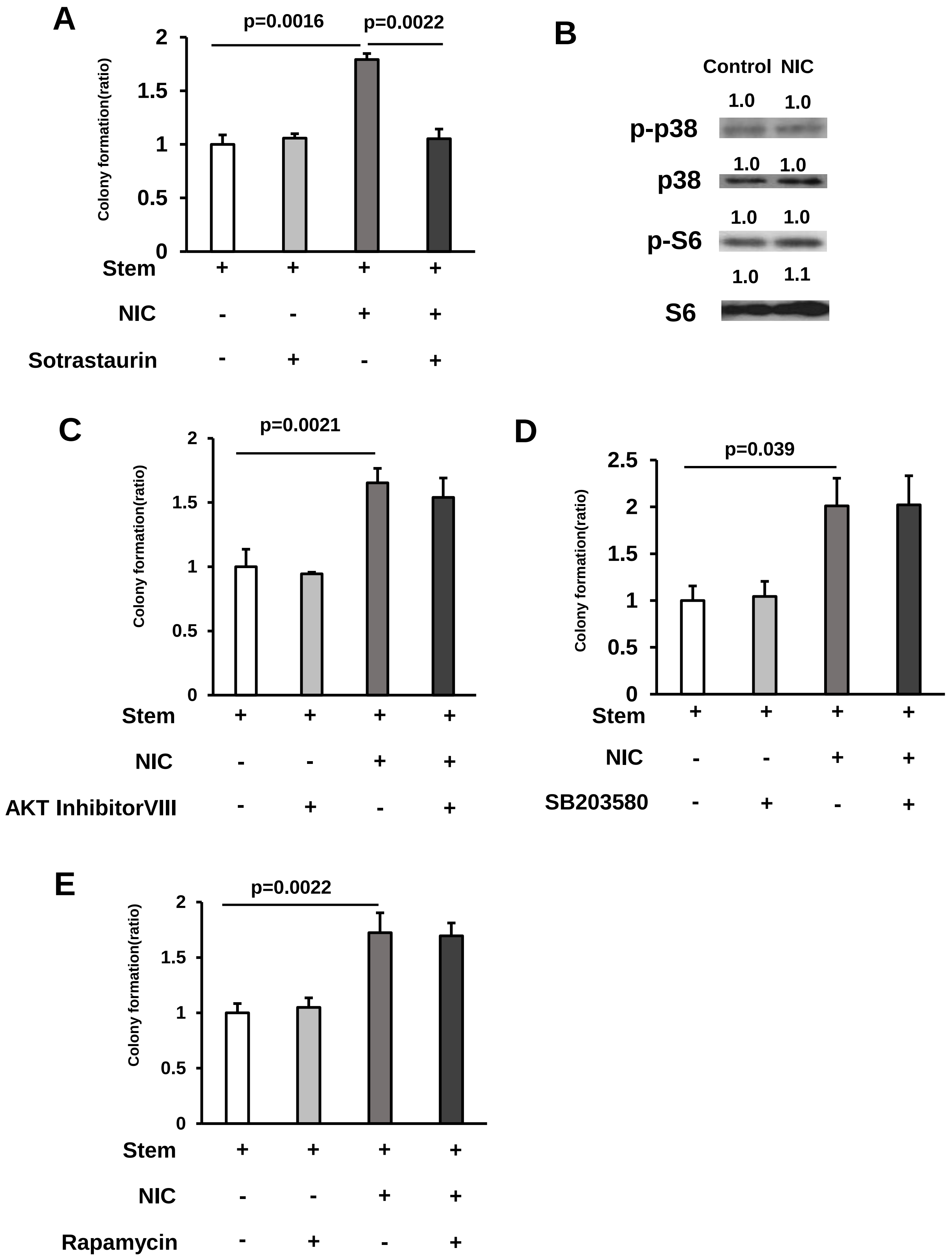


### Figure3 Supplemental

A

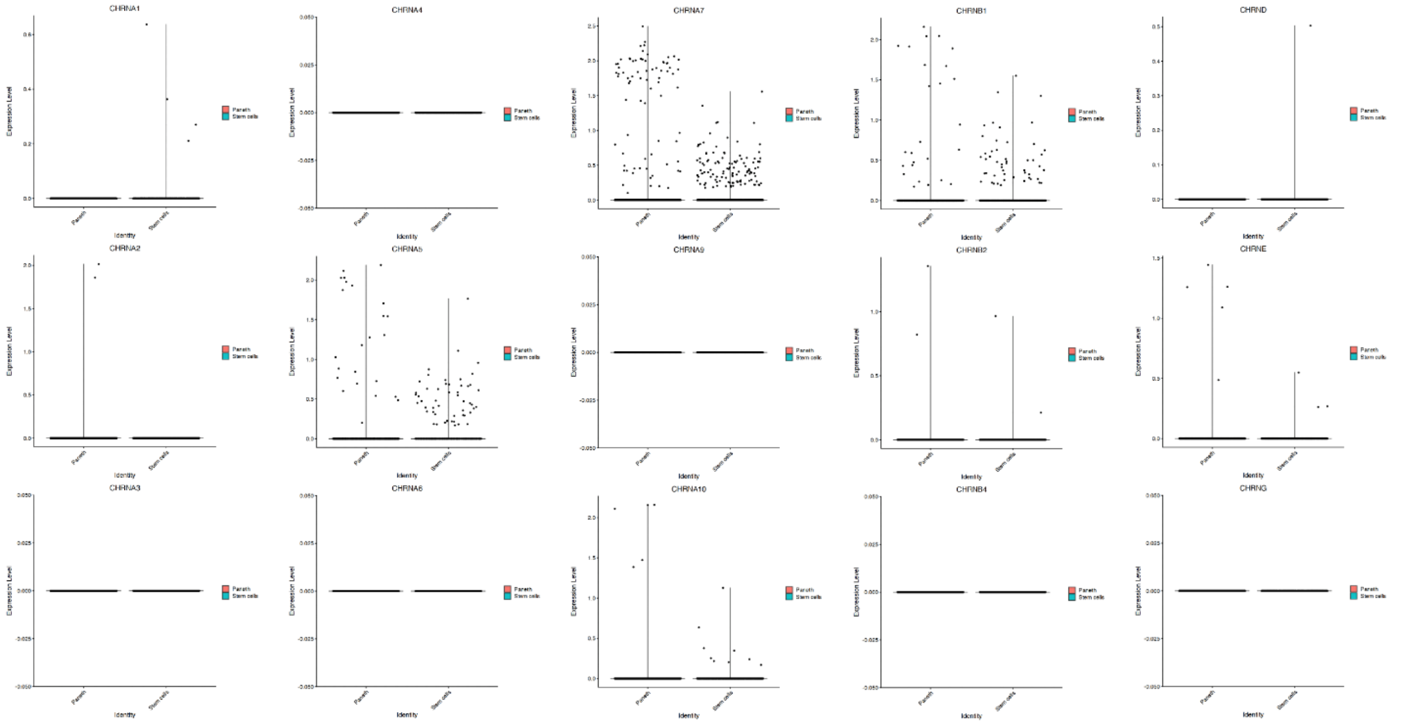

B

| | Paneth cells | Stem cells |
|---|---|---|
| All | 2655 | 1022 |
| CHRNA1 | 0 | 4 |
| CHRNA2 | 2 | 0 |
| CHRNA3 | 0 | 0 |
| CHRNA4 | 0 | 0 |
| CHRNA5 | 24 | 52 |
| CHRNA6 | 0 | 0 |
| CHRNA7 | 78 | 116 |
| CHRNA8 | 0 | 0 |
| CHRNA9 | 0 | 0 |
| CHRNA10 | 5 | 9 |
| CHRNB1 | 26 | 51 |
| CHRNB2 | 2 | 2 |
| CHRNB3 | 0 | 0 |
| CHRNB4 | 0 | 0 |
| CHRND | 0 | 1 |
| CHRNE | 5 | 3 |
| CHRNG | 0 | 0 |

### Figure4 Supplemental

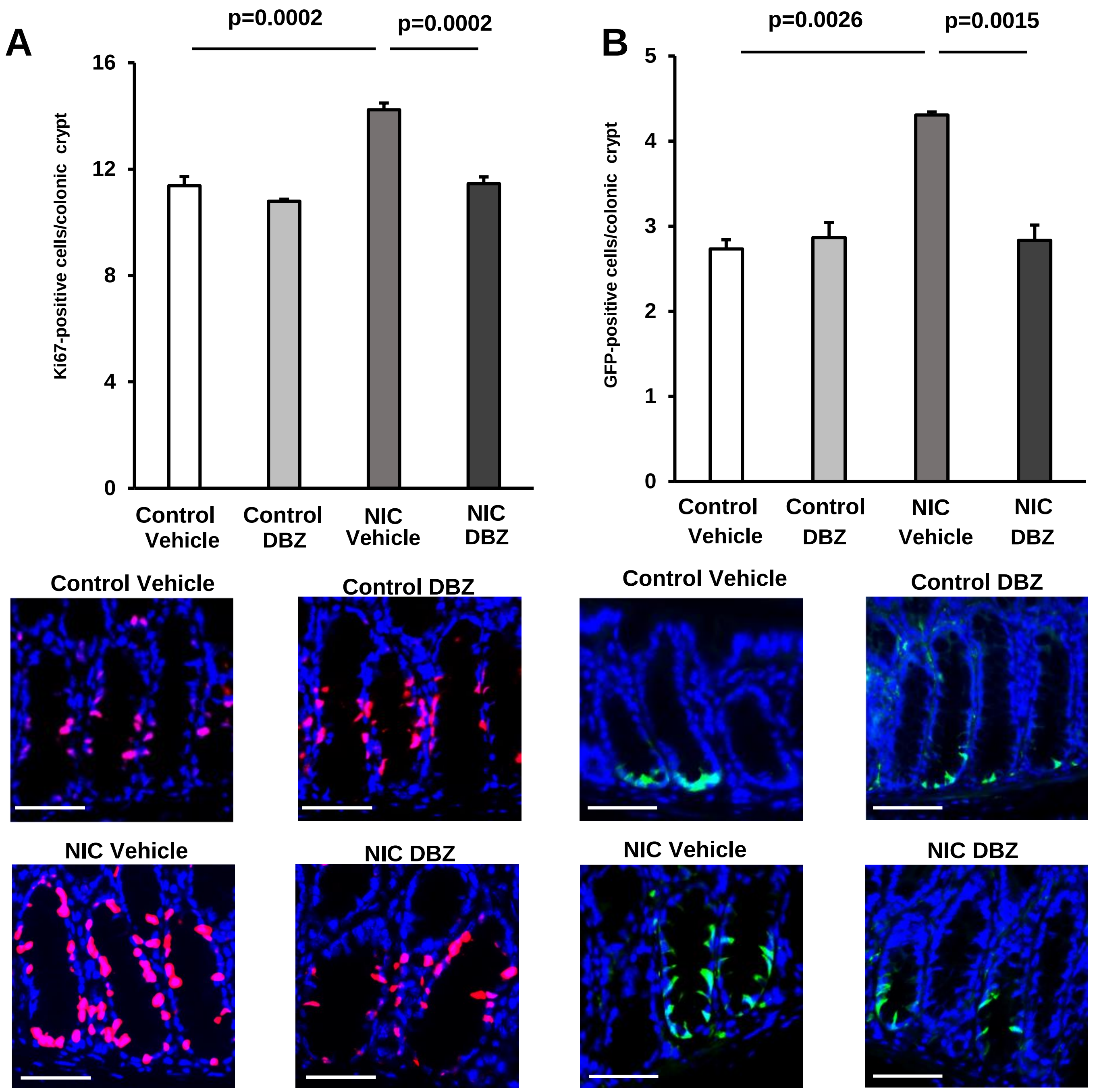


### Figure5 Supplemental
