## Supplemental Table1 for "Nicotine enhances the stemness and tumorigenicity in intestinal stem cells via Hippo-YAP/TAZ and Notch signal pathway"

**Primer sequences used in qRT-PCR**

| | **Forward primer** | **Reverse Primer** |
|---|---|---|
| $\alpha$1 | TATAACAACGCAGACGGCGA | CACAGAGACCGTCATAGGTCC |
| $\alpha$2 | GTGCCCAACACTTCCGATG | TGTAGTCATTCCATTCCTGCTTT |
| $\alpha$3 | CCAGTTTGAGGTGTCTATGTC | TCGGCGTTGTTGTAAAGC |
| $\alpha$4 | CTCAGATGTGGTCCTTGTC | GAGTTCAGATGGGATGCG |
| $\alpha$5 | CATCGTTTTGTTTGATAATGC | TGCGTCCAAGTGACAGTG |
| $\alpha$6 | TGTCTCCGATCCCGTCAC | TTGTTATACAGAACGATGTCAGG |
| $\alpha$7 | GGTCATTTGCCCACTCTG | GACAGCCTATCGGGTGAG |
| $\alpha$9 | ACAAGGCCACCAACTCCA | ACCAACCCACTCCTCCTCTT |
| $\alpha$10 | TCTGACCTCACAACCCACAA | TCCTGTCTCAGCCTCCATGT |
| $\beta$1 | AAGCCGAAGGCCAACTGATTA | TCCTGCCTCTCCTCTCCTTC |
| $\beta$2 | CCGGCAAGAAGCCGGGACCT | CTCGCTGACACAAGGGCTGCG |
| $\beta$3 | AAGAAGCAGACTCCTACC | AACAACCTGACTGATGAAG |
| $\beta$4 | CTACAGGAAGCATTAGAGG | CAGAATACACACAATCACG |
| Hes1 | ACACCGGACAAACCAAAGAC | AATGCCGGGAGCTATCTTTC |
| Hes5 | GCAGCATAGAGCAGCTGAAG | AGGCTTTGCTGTGTTTCAGG |
| HeyL | GTCTTGCAGATGACCGTGGA | CTCGGGCATCAAAGAACCCT |
| Hey1 | CACCTGAAAATGCTGCACAC | ATGCTCAGATAACGGGCAAC |
| Hes1 | ACACCGGACAAACCAAAGAC | AATGCCGGGAGCTATCTTTC |
| Jagged1 | CCTCGGGTCAGTTTGAGCTG | CCTTGAGGCACACTTTGAAGTA |
| Jagged2 | ACGAGGAGGATGAAGAGCTGA | GGGGTCTTTGGTGAACTTGTG |
| YAP | CGCTCTTCAATGCCGTCATG | AGTCATGGCTTGCTCCCATC |
| TAZ | TCTGTCATGAACCCCAAGCC | GGTGGTTCTGTGGACTCAGG |
| 18S | GTAACCCGTTGAACCCCATT | CCATCCAATCGGTAGTAGCG |
